# Curated compendium of human transcriptional biomarker data

**DOI:** 10.1101/191064

**Authors:** Nathan P. Golightly, Anna I. Bischoff, Avery Bell, Parker D. Hollingsworth, Stephen R. Piccolo

## Abstract

Genome-wide transcriptional profiles provide broad insights into cellular activity. One important use of such data is to identify relationships between transcription levels and patient outcomes. These translational insights can guide the development of biomarkers for predicting outcomes in clinical settings. Over the past decades, data from many translational-biomarker studies have been deposited in public repositories, enabling other scientists to reuse the data in follow-up studies. However, data-reuse efforts require considerable time and expertise because transcriptional data are generated using heterogeneous profiling technologies, preprocessed using diverse normalization procedures, and annotated in non-standard ways. To address this problem, we curated a compendium of 45 translational-biomarker datasets from the public domain. To increase the data’s utility, we reprocessed the raw expression data using a standard computational pipeline and standardized the clinical annotations in a fully reproducible manner (see osf.io/ssk3t). We believe these data will be particularly useful to researchers seeking to validate gene-level findings or to perform benchmarking studies—for example, to compare and optimize machine-learning algorithms’ ability to predict biomedical outcomes.

## Background&summary

DNA encodes a cell’s instruction manual in the form of genes and regulatory sequences^1^. Cells behave differently, in part, because genes are transcribed into RNA at different levels within those cells^2^. Researchers examine gene-expression levels to understand cellular dynamics and the mechanisms behind cellular aberrations, including those that lead to disease development. Modern technologies now make it possible to profile expression levels for thousands of genes at a time for a modest expense^3^. Using these high-throughput technologies, scientists have performed thousands of studies to characterize biological processes and to evaluate the potential for precision-medicine applications. One such application is to derive *transcriptional biomarkers*—patterns of expression that indicate disease states or that predict medical outcomes, such as relapse, survival, or treatment response^4–10^. Indeed, already to date, more than 100 transcriptional biomarkers have been proposed for predicting breast-cancer survival alone^11^.

Many funding agencies and academic journals have imposed policies that require scientists to deposit transcriptional data in publicly accessible databases. These policies seek to ensure that other scientists can verify the original study’s findings and can reuse the data in secondary analyses. For example, Gene Expression Omnibus (GEO) currently contains data for more than 2 million biological samples^12^. Upon considering infrastructure and personnel costs, we estimate that these data represent hundreds of millions—if not billions—of dollars (USD) of collective research investment. Reusing these vast resources offers an opportunity to reap a greater return on investment—perhaps most importantly via informing and validating new studies. Unfortunately, although anyone can access GEO data, researchers vastly underutilize this treasure trove because preparing data for new analyses requires considerable background knowledge and informatics expertise.

In GEO, data are typically available in two forms: 1) raw data, as produced originally by the data-generating technology, and 2) processed data, which were used in the data generators’ analyses. In most cases, researchers process raw data in a series of steps that might include quality-control filtering, noise reduction, standardization, and summarization (e.g., summarizing to gene-level values and excluding outliers). Data from different profiling technologies must be handled in ways that are specific to each technology. However, even for datasets generated using the same profiling technology, the methods employed for data preprocessing vary widely across studies. This heterogeneity makes it difficult for researchers to perform secondary analyses and to trust that analytical findings are driven primarily by biological mechanisms rather than differences in data preprocessing. In addition, when data have not been mapped to biologically meaningful identifiers, it may be difficult for researchers to draw biological conclusions from the data.

Sample-level annotations accompany each GEO dataset. For biomarker studies, such metadata might include medical diagnoses or treatment outcomes, as well as covariates such as age, sex, or ethnicity. Although GEO publishes metadata in a semi-standardized format and bioinformatics tools exist for downloading and parsing GEO data^13,14^, it is difficult for many researchers to extract these data into a form that is suitable for secondary analyses. Within annotation files, values are often stored in key/value pairs with nondescript column names. Many columns are not useful for analytical purposes (e.g., when all samples have the same value). When values are missing, the columns often become shifted; accordingly, data for a given variable may be spread across multiple columns. Moreover, a variety of descriptors (e.g., “?”, “N/A”, or “Unknown”) are used to indicate missing values,thus requiring the analyst to account for these differences. In addition, seemingly minor errors, such as spelling mistakes or inconsistent capitalization, can hamper secondary-analysis efforts.

In response to these challenges, we compiled the *Biomarker Benchmark*, a curated compendium of 45 transcriptional-biomarker datasets from GEO. These datasets represent a variety of human-disease states and outcomes, many related to cancer. We obtained raw gene-expression files, renormalized them using a common algorithm, and summarized the data using gene-level annotations (Figure 1). We used two techniques to check for quality-control issues in the gene-expression data. For datasets where gene-expression data were processed in multiple batches—and where batch information was available—we corrected for batch effects. Finally, we prepared a version of the data that is suitable for direct application in machine-learning analyses. We standardized continuous values, one-hot encoded discrete values, and imputed missing values.

**Fig. 1.**
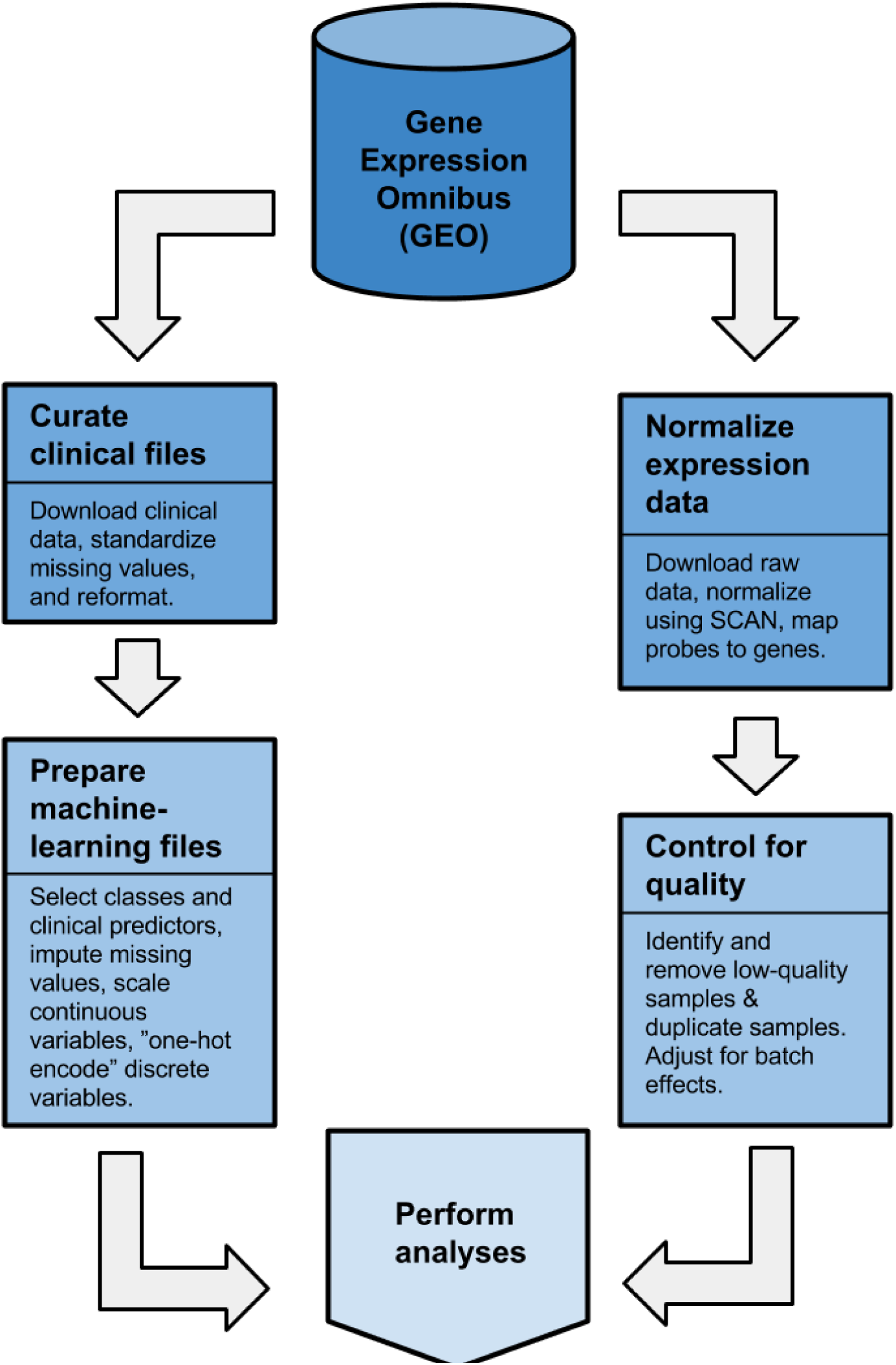
Flow diagram that illustrates the process we used to collect and curate the data. We wrote computer scripts that downloaded the data, checked for quality, normalized and standardized data values, and stored the data in analysis-ready file formats. The specific steps differed for clinical and expression data (see Methods).

## Methods

### Selecting data

To select datasets, we executed a custom search in Gene Expression Omnibus (GEO). First, we limited our search to data series that were associated with the Medical Subject Heading (MeSH) term "biomarker" and that came from *Homo sapiens* subjects. Next we limited the search to data generated using Affymetrix gene-expression microarrays and for which raw expression data were available (so we could renormalize the data). For each dataset, we examined the metadata to ensure that each series had at least one biomarker-relevant clinical variable. These included variables such as prognosis, disease stage, histology, and treatment success or relapse. Lastly, we selected series that included data for at least 70 samples (before additional filtering, see below).

Based on these criteria, we identified 36 GEO series. Two series (GSE6532 and GSE26682) contained data for two types of Affymetrix microarray. To avoid platform-related biases, we separated each of these series into two datasets; we useda suffix for each that indicates the microarray platform (e.g., GSE6532_U133A and GSE6532_U133Plus2). For both of these datasets, the biological samples profiled using either microarray platform were distinct. The GSE2109 series—known as the Expression Project for Oncology (expO)—had been produced by the International Genomics Consortium and contains data for 129 different cancer types. To avoid confounding effects due to tissue-specific expression and because the metadata differed considerably across the cancer types, we split GSE2109 into multiple datasets based on cancer type (Table 1). We excluded tissue types for which fewer than 70 samples were available; we also excluded the “omentum” cancer type because it was relatively heterogeneous and had relatively few samples.

We used publicly available data for this study and played no role in contacting the research subjects. We received approval to work with these data from Brigham Young University’s Institutional Review Board (E 14522).

**Table 1.** Overview of data sources used in this study.

| Series ID<sup>a</sup> | Tissue type(s) | # Samples | Clinical variable(s) | # Genes | Affymetrix Platform(s) |
| --- | --- | --- | --- | --- | --- |
| GSE1456<sup>25</sup> | Breast cancer | 157 | Elston grade; overall survival status; overall survival time; relapse status; relapse time | 11832 | Genome U133A |
| GSE2109<sup>26</sup> | Breast | 263 | Age; alcohol consumption; days from diagnosis to excision; ER status; ethnic background; ethnic background; family history of cancer; fibrocystic disease; Her2 status; histology; hormonal therapy duration; mammogram status and findings; metastasis; metastatic sites; multiple tumors; node involvement; oophorectomy status; oral contraceptive use; PR status; prior therapy status; quality metric; relapse time; retreatment states; sex; stage; tobacco use; tumor grade; tumor size | 20024 | Genome U133 Plus 2.0 |
| GSE2109<sup>26</sup> | Colon | 255 | Age; alcohol consumption; days from diagnosis to excision; diagnosis method; Dukes' stage; ethnic background; family history of cancer; histology; metastasis; metastatic sites; multiple tumors; node involvement; primary site; prior screening status; prior therapy status; quality metric; relapse time; retreatment states; sex; stage; symptoms; tobacco use; tumor grade; tumor size | 20024 | Genome U133 Plus 2.0 |
| GSE2109<sup>26</sup> | Endometrium | 51 | Age; alcohol consumption; days from diagnosis to excision; ethnic background; family history of cancer; histology; metastasis; metastatic sites; multiple tumors; node involvement; primary site; quality metric; stage; symptoms; tobacco use; tumor grade; tumor size | 20024 | Genome U133 Plus 2.0 |
| GSE2109<sup>26</sup> | Kidney | 209 | Age; alcohol consumption; days from diagnosis to excision; ethnic background; family history of cancer; histology; metastasis; metastatic sites; multiple tumors; primary site; prior therapy status; quality metric; relapse time; retreatment states; sex; stage; tobacco use; tumor grade; tumor size | 20024 | Genome U133 Plus 2.0 |
| GSE2109<sup>26</sup> | Lung | 103 | Age; alcohol consumption; days from diagnosis to excision; ethnic background; family history of cancer; histology; metastasis; metastatic sites; multiple tumors; node involvement; primary site; prior therapy status; quality metric; relapse time; retreatment states; sex; stage; symptoms; tobacco use; tumor grade; tumor size | 20024 | Genome U133 Plus 2.0 |
| GSE2109<sup>26</sup> | Ovary | 158 | Age; alcohol consumption; days from diagnosis to excision; esophagitis reflux history; ethnic background; family history of cancer; fibrocystic disease; histology; mammogram history; metastasis; metastatic sites; multiple tumors; node involvement; node involvement; oophorectomy status; primary site; prior therapy status; quality metric; relapse time; retreatment states; screening history; stage; symptoms; tobacco use; tumor grade; tumor size | 20024 | Genome U133 Plus 2.0 |
| GSE2109<sup>26</sup> | Prostate | 79 | Age; alcohol consumption; days from diagnosis to excision; diagnosis method; ethnic background; family history of cancer; Gleason score; histology; metastasis; multiple tumors; node involvement; prior therapy status; prostate-specific antigen (PSA) testing history; PSA finding; quality metric; stage; symptoms; tobacco use; tobacco use; tumor grade; tumor size | 20024 | Genome U133 Plus 2.0 |
| GSE2109<sup>26</sup> | Uterine | 112 | Age; alcohol consumption; days from diagnosis to excision; ethnic background; family history of cancer; histology; human papilloma virus diagnosis history; metastasis; metastatic sites; multiple tumors; node involvement; primary site; prior therapy status; quality metric; relapse; stage; symptoms; tobacco use; tumor grade; tumor size | 20024 | Genome U133 Plus 2.0 |
| GSE4271<sup>27,28</sup> | Glial | 100 | Age; recurrence status; sex; survival status; survival time; WHO grade | 11832 | Genome U133A |
| GSE5460<sup>29</sup> | Breast cancer | 127 | ER status; Her2 status; histological type; lymphovascular invasion; node status; tumor grade; tumor size | 20024 | Genome U133 Plus 2.0 |
| GSE5462<sup>30,31</sup> | Breast cancer | 52 | Treatment history; treatment response | 11832 | Genome U133A |
| GSE6532<sup>32</sup> | Breast carcinoma | 317 | Age; distant metastasis-free survival time/status; ER status; genomic grade index; node involvement; PR status; recurrence-free survival time/status; tumor grade; tumor size | 11832 | Genome U133A |
| GSE6532<sup>32</sup> | Breast carcinoma | 87 | Age; distant metastasis-free survival time/status; ER status; genomic grade index; node involvement; PR status; recurrence-free survival time/status; tumor grade; tumor size | 20024 | Genome U133 Plus 2.0 |
| GSE10320<sup>33</sup> | Wilms Tumor | 144 | Relapse | 11832 | Genome U133A |
| GSE15296<sup>34</sup> | Peripheral Blood | 75 | Kidney transplant rejection; subtype | 20024 | Genome U133 Plus 2.0 |
| GSE19804<sup>35</sup> | Paired tumor and normal tissues | 60 | Age; tissue type; tumor stage | 20024 | Genome U133 Plus 2.0 |
| GSE20181<sup>31,36</sup> | Breast cancer | 50 | Treatment history; treatment response | 11832 | Genome U133A |
| GSE20189<sup>37</sup> | Lung adenocarcinoma | 162 | Case/control status; morphology; smoking status; stage | 11832 | Genome U133A 2.0 |
| GSE21510<sup>38</sup> | Laser capture microdissection and homogenized tissues (surgically resected material) | 104 | Metastasis; stage; tissue type | 20024 | Genome U133 Plus 2.0 |
| GSE25507<sup>39</sup> | Peripheral blood lymphocyte | 146 | Case/control status (autism); paternal age, maternal age, subject age | 20024 | Genome U133 Plus 2.0 |
| GSE26682<sup>40-42</sup> | Colorectal tumor | 140 | Age; microsatellite instability status; sex | 11832 | Genome U133A |
| GSE26682<sup>40-42</sup> | Colorectal tumor | 160 | Age; microsatellite instability status; sex | 20024 | Genome U133 Plus 2.0 |
| GSE27279<sup>43</sup> | Posterior Fossa Ependymoma | 100 | Age; sex; tumor location | 16632 | Exon 1.0 ST |
| GSE27342<sup>44,45</sup> | Paired gastric tumor and normal tissue | 72 | Age; sex; stage; tissue type; tumor grade | 16632 | Exon 1.0 ST |
| GSE27854<sup>46</sup> | Colorectal tumor | 115 | Metastasis; stage | 20024 | Genome U133 Plus 2.0 |
| GSE30219<sup>47</sup> | Lung | 293 | Age; follow-up time; histology; metastasis; node involvement; relapse status; sex; survival; survival time; tumor size | 20024 | Genome U133 Plus 2.0 |
| GSE30784<sup>48</sup> | Oral squamous cell carcinoma | 229 | Age; case/control status; sex | 20024 | Genome U133 Plus 2.0 |
| GSE32646<sup>49</sup> | Breast | 115 | Age; ER status (IHC); Her2 status (FISH); histological grade; lymph node involvement; pathologic complete response; PR status (IHC); stage; tumor size | 20024 | Genome U133 Plus 2.0 |
| GSE37147<sup>50</sup> | Bronchial sample | 238 | Age; case/control status (COPD); FEV1/FVC score/percentage; inhaled medication status; sex; smoking status; tobacco use | 21614 | Gene 1.0 ST |
| GSE37199<sup>51</sup> | Blood sample | 94 | Disease stage (advanced castration resistant prostate cancer) | 20024 | Genome U133 Plus 2.0 |
| GSE37745<sup>52</sup> | Non-small cell lung cancer | 196 | Adjuvant treatment status; age; histology; recurrence time/status; sex; stage; survival time/status; WHO performance status | 20024 | Genome U133 Plus 2.0 |
| GSE37892<sup>53</sup> | Stage-II colon carcinoma | 130 | Age; diagnosis history; localisation; stage; time until metastasis | 20024 | Genome U133 Plus 2.0 |
| GSE38958<sup>54</sup> | Peripheral blood mononuclear cell | 115 | Age; diagnosis (Idiopathic pulmonary fibrosis); ethnicity; predicted FVC percent; sex | 16632 | Exon 1.0 ST |
| GSE39491<sup>55</sup> | Esophageal and gastric samples | 120 | Tumor cell type | 11832 | Genome U133 Plus 2.0 |
| GSE39582<sup>56</sup> | Colon cancer | 566 | Adjuvant chemotherapy; age; BRAF mutation status; chromosome instability status; CIMP status; KRAS mutation status; mismatch repair status; overall survival time/status; recurrence-free survival time/status; sex; stage; TP53 mutation status; tumor location | 20024 | Genome U133 Plus 2.0 |
| GSE40292<sup>57</sup> | Afferent limb tissue and whole-blood sample | 195 | Diagnosis; sex | 21614 | Gene 1.0 ST |
| GSE43176<sup>58</sup> | Leukemic blast sample | 104 | Cytogenetics; disease state; FAB stage; KRAS mutation status; NRAS mutation status; subtype | 11832 | Genome U133A |
| GSE46449<sup>59</sup> | Peripheral blood leukocyte | 53 | Age; diagnosis (bipolar disorder) | 20024 | Genome U133 Plus 2.0 |
| GSE46691<sup>60</sup> | Prostate | 545 | Gleason score; metastasis | 16632 | Exon 1.0 ST |
| GSE46995<sup>61</sup> | Leukocyte | 85 | Age; disease status (biliary atresia) | 21614 | Gene 1.0 ST |
| GSE48391<sup>62</sup> | Breast | 81 | ER status; gene-expression subtype; Her2 status; recurrence status; survival time/status | 20024 | Genome U133 Plus 2.0 |
| GSE58697<sup>63</sup> | Desmoid tumor | 72 | Age; follow-up time; recurrence time; sex; tumor location; tumor size | 20024 | Genome U133 Plus 2.0 |
| GSE63885<sup>64</sup> | Ovarian cancer surgical sample | 101 | Adjuvant chemotherapy; BRCA mutation status; clinical status at last follow-up, clinical status after 1st line chemotherapy; disease-free survival; FIGO stage; histopathological type; overall survival; residual tumor size; TP53 accumulation in cancer cells (IHC); TP53 mutation status; TP53 mutation status; tumor grade | 20024 | Genome U133 Plus 2.0 |
| GSE67784<sup>65</sup> | Peripheral blood sample | 309 | Sex; V30M mutation status; whether exhibiting symptoms | 21614 | Gene 1.1 ST |

<sup>a</sup> These identifiers represent data series in Gene Expression Omnibus. Some identifiers are listed multiple times; in these cases, we used a subset of the series data (for a specific tissue type or microarray platform).

### Preparing clinical annotations

For each dataset, we wrote custom R scripts ^15^ that download, parse, and reformat the clinical annotations. Initially, these scripts download data using the *GEOquery* package ^13^. Next they generate a tab-delimited text file for each dataset that contains all available clinical annotations, except those with identical values for all samples (for example, platform name, species name, submission date) or that were unique to each biological sample (for example, sample title). In addition, these scripts generate Markdown files daringfireball.net/projects/markdown/syntax that summarize each dataset and indicate sources.

In some cases, multiple data values are included in the same cell in GEO annotation files. For example, in GSE5462, one patient’s clinical demographics and treatment responses are listed as "female;breast tumor;Letrozole, 2.5mg/day,oral,10-14 days; responder." We parsed these values and split them into separate columns for each sample. After these cleaning steps, the datasets contained 7.8 variables of metadata, on average ( Table 1). Next we searched each dataset for missing values. Across the datasets, 11 distinct expressions had been used by the original data generators to represent missingness; these included "N/A", "NA","MISSING", "NOT AVAILABLE", "?", and others. To support consistency, we standardized these values across the datasets, using a value of "NA". On average, 17.0% of the metadata values were missing per dataset; this proportion differed considerably across the datasets (Figure 2).

**Fig. 2.**
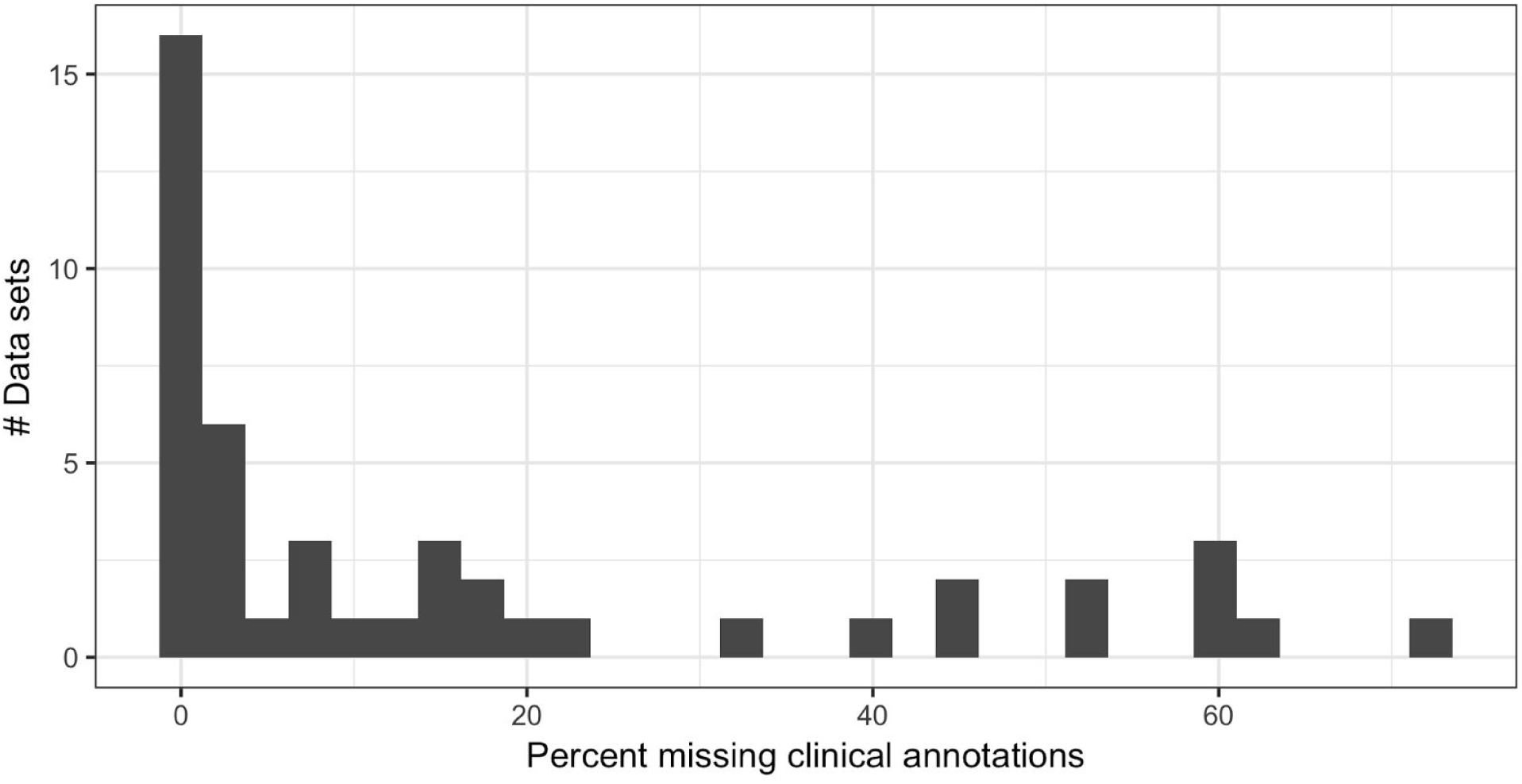
Histogram showing the proportion of missing clinical-annotation values per dataset. Some datasets contained no missing values, while others were missing as many as as 72.3% of data values.

We anticipate that many researchers will use these data to develop and benchmark machine-learning algorithms. Accordingly, we have prepared secondary versions of the clinical annotations that are ready to use in machine-learning analyses. First, we identified class variables that have potential relevance for biomarker applications. In many cases, these variables were identical to those used in the original studies; we also included class variables that had not been used in the original studies. On average, the datasets contain 2.9 class variables. Second, we identified clinical variables that could be used as predictor variables (or covariates). Using these data, we generated one output file per class variable or predictor variable and named the output files using descriptive prefixes (e.g., "Prognosis", "Diagnosis", or "Stage"). The same variable can be used as a class variable in one context and a predictor in a different context. When a given sample was missing data for a given class variable, we excluded that sample from the respective output file for that class variable. After this filtering step, we identified class variables with fewer than 40 samples and excluded these class variables. When predictor variables were missing more than 20% data (Figure 2), we did not generate an output file for these variables. When predictor variables were missing less than 20% data, we imputed missing values using median-based imputation for continuous variables and mode-based imputation for discrete variables^16^. We scaled continuous predictor variables to have zero mean and unit variance. We transformed discrete predictor variables using one-hot encoding; each unique value, except the first, was treated as a binary variable. In cases where discrete values were rare, we merged values. For example, in GSE2109_Breast, we merged *Pathological_Stage* values 3A, 3B, 3C, and 4 into a category called “3-4” because relatively few patients fell into the individual categories (38, 8, 22, and 5 samples, respectively). In addition, some class variables were ordinal in nature (e.g., cancer stage or tumor grade); we transformed these to binary variables. Finally, some clinical outcomes were survival or relapse times; we transformed these data to (discrete) class variables, using a threshold to distinguish between "long-term" and "short-term" survivors and excluding patients who were censored after the survival threshold had been reached. Our computer scripts (see *Code availability*) encode these decisions for each dataset.

### Preprocessing gene-expression data

We created a computational pipeline (using R and shell scripts) that downloads, normalizes, and standardizes the raw-expression data. We used the *GEOquery* package^13^ to download the CEL files and then normalized them using the *SCAN.UPC* package<u>^17^</u>. Some heterogeneity exists even among platforms from the same manufacturer (Affymetrix). The number of probes and the probe sequences used in designing the microarray architectures vary. To help mitigate this heterogeneity and to aid in biological interpretation, we summarized the data using gene-level annotations from *Brainarray*^18^

### Code availability

Our computer scripts are stored in the open-access *Biomarker Benchmark* repository (osf.io/ssk3t). Accordingly, other researchers can reproduce our curation process and produce alternative versions of the data.

## Data records

After filtering (see Methods), we collected data for 7,037 biological samples across 45 datasets (Table 1). On average, the data sets contain values for 18,043 genes (Table 1). In total, our repository contains 129 class variables (2.8 per dataset) and 2.1 unique values per class variable.

All output data are stored in tab-delimited text files and are structured using the “tidy data” methodology^19^. Accordingly, data users can import the files directly into analytical tools such as Microsoft Excel, R, or Python. All data files are publicly and freely available in the open-access *Biomarker Benchmark* repository (osf.io/ssk3t">). The original data files are available via Gene Expression Omnibus using the accession numbers listed in Table 1.

## Technical validation

We evaluated each sample using the *IQRray*^20^ software, which produces a quality score for individual samples. Using these metrics, we applied Grubb’s statistical test (*outliers* package^21^) to each dataset, identified poor-quality outliers (Figure 3), and excluded these samples (Table 2). Next we used the *DoppelgangR* package^22^ to identify samples that may have been duplicated inadvertently. We manually reviewed sample pairs that *DoppelgangR* flagged as potential duplicates. We excluded most sample pairs that were flagged (Table 2), even if the clinical annotations for both samples were distinct, under the assumption that these samples had somehow been mislabeled. In GSE46449, many samples were biological replicates, and we retained one of each replicate set. GSE5462, GSE19804, and GSE20181 contained samples that had been profiled in a paired manner (e.g., pre-and post-treatment); we retained these samples.

**Fig. 3.**
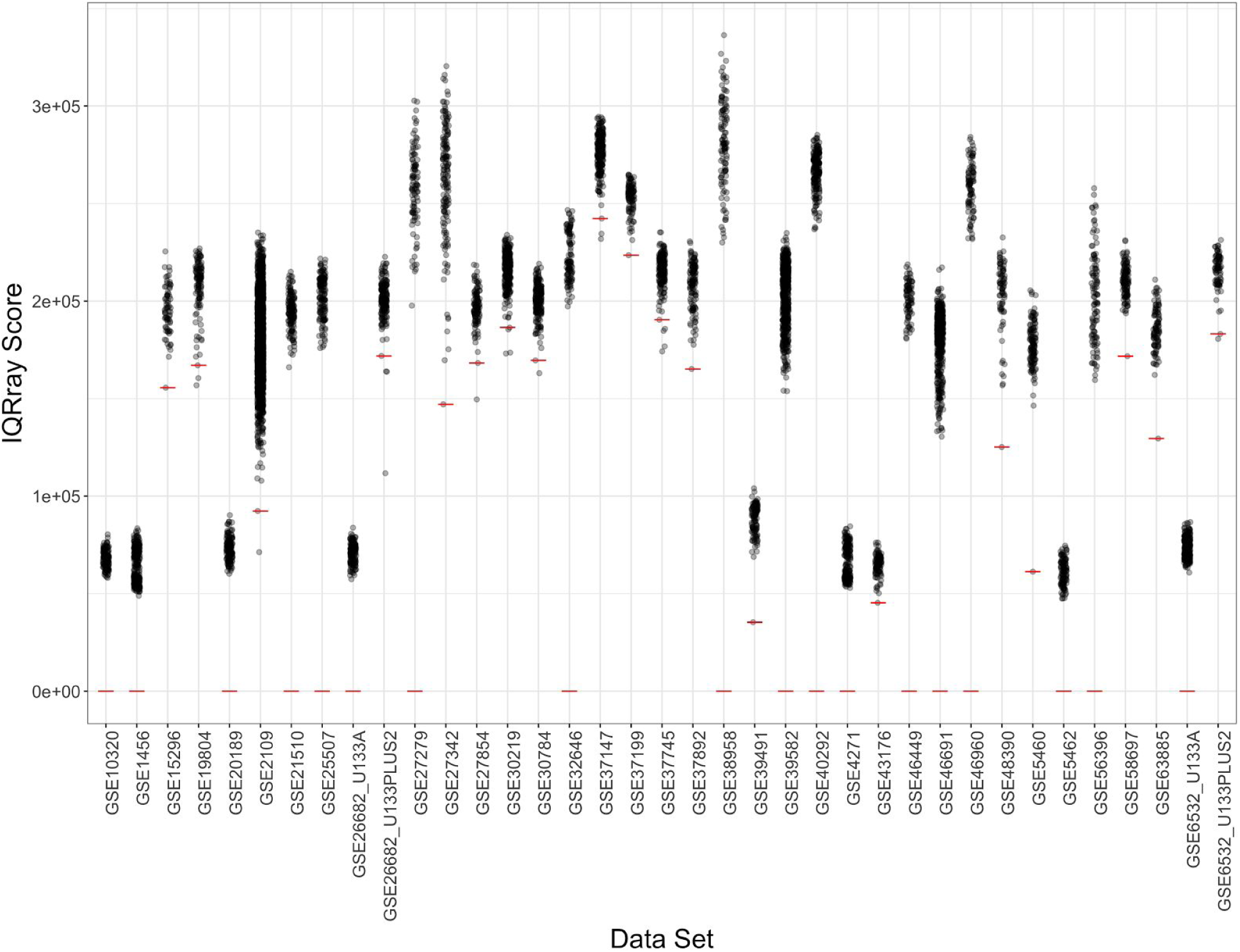
Distribution of IQRay quality scores for each dataset. Sample qualities are plotted for each dataset. Low-quality samples were identified using Grubb’s test. Samples that fall on or below the red threshold were excluded from the data repository.

**Table 2.** Summary of excluded samples. We excluded samples that did not pass our quality-control criteria or that appeared to be duplicated. The Gene Expression Omnibus seriesand sample identifiers are listed, along with the reason we excluded each sample.

| Series | Sample | Reason |
| --- | --- | --- |
| GSE15296 | GSM382283 | Poor Quality |
| GSE19804 | GSM494596 | Poor Quality |
| GSE19804 | GSM494654 | Poor Quality |
| GSE19804 | GSM494657 | Poor Quality |
| GSE20181 | GSM506289 | Duplicate |
| GSE20181 | GSM506294 | Duplicate |
| GSE20181 | GSM506304 | Duplicate |
| GSE20181 | GSM125198 | Duplicate |
| GSE20181 | GSM125210 | Duplicate |
| GSE20181 | GSM125230 | Duplicate |
| GSE2109_Breast | GSM53059 | Duplicate |
| GSE2109_Breast | GSM53027 | Duplicate |
| GSE2109_Colon | GSM89040 | Duplicate |
| GSE2109_Colon | GSM152664 | Duplicate |
| GSE2109_Colon | GSM152632 | Duplicate |
| GSE2109_Colon | GSM179922 | Duplicate |
| GSE2109_Colon | GSM89044 | Duplicate |
| GSE2109_Colon | GSM152666 | Duplicate |
| GSE2109_Colon | GSM179820 | Duplicate |
| GSE2109_Colon | GSM179924 | Duplicate |
| GSE2109_Lung | GSM203652 | Poor Quality |
| GSE2109_Ovary | GSM76554 | Duplicate |
| GSE2109_Ovary | GSM203725 | Duplicate |
| GSE2109_Ovary | GSM76567 | Duplicate |
| GSE2109_Ovary | GSM231913 | Duplicate |
| GSE2109_Ovary | GSM46839 | Poor Quality |
| GSE2109_Prostate | GSM179790 | Duplicate |
| GSE2109_Prostate | GSM179843 | Duplicate |
| GSE2109_Prostate | GSM179903 | Duplicate |
| GSE25507 | GSM627091 | Duplicate |
| GSE25507 | GSM627087 | Duplicate |
| GSE25507 | GSM627096 | Duplicate |
| GSE25507 | GSM627078 | Duplicate |
| GSE25507 | GSM627085 | Duplicate |
| GSE25507 | GSM627196 | Duplicate |
| GSE25507 | GSM627153 | Duplicate |
| GSE25507 | GSM627180 | Duplicate |
| GSE25507 | GSM627099 | Duplicate |
| GSE25507 | GSM627115 | Duplicate |
| GSE25507 | GSM627118 | Duplicate |
| GSE25507 | GSM627124 | Duplicate |
| GSE25507 | GSM627154 | Duplicate |
| GSE25507 | GSM627204 | Duplicate |
| GSE25507 | GSM627209 | Duplicate |
| GSE25507 | GSM627215 | Duplicate |
| GSE26682 | GSM656833 | Duplicate |
| GSE26682 | GSM656770 | Duplicate |
| GSE26682_U133PLUS2 | GSM656860 | Poor Quality |
| GSE26682_U133PLUS2 | GSM656613 | Poor Quality |
| GSE26682_U133PLUS2 | GSM656839 | Poor Quality |
| GSE26682_U133PLUS2 | GSM656721 | Poor Quality |
| GSE27342 | GSM675945 | Duplicate |
| GSE27342 | GSM675947 | Duplicate |
| GSE27342 | GSM675933 | Duplicate |
| GSE27342 | GSM675935 | Duplicate |
| GSE27342 | GSM676040 | Poor Quality |
| GSE27342 | GSM687519 | Poor Quality |
| GSE27854 | GSM687525 | Poor Quality |
| GSE30219 | GSM748210 | Duplicate |
| GSE30219 | GSM748212 | Duplicate |
| GSE30219 | GSM748218 | Duplicate |
| GSE30219 | GSM748219 | Duplicate |
| GSE30219 | GSM748255 | Poor Quality |
| GSE30219 | GSM748247 | Poor Quality |
| GSE30219 | GSM748057 | Poor Quality |
| GSE30219 | GSM748266 | Poor Quality |
| GSE30784 | GSM764928 | Duplicate |
| GSE30784 | GSM764930 | Duplicate |
| GSE30784 | GSM764904 | Poor Quality |
| GSE30784 | GSM764970 | Poor Quality |
| GSE32646 | GSM809214 | Duplicate |
| GSE32646 | GSM809248 | Duplicate |
| GSE32646 | GSM809251 | Duplicate |
| GSE32646 | GSM809254 | Duplicate |
| GSE37147 | GSM912230 | Duplicate |
| GSE37147 | GSM912296 | Duplicate |
| GSE37147 | GSM912296 | Duplicate |
| GSE37147 | GSM912305 | Duplicate |
| GSE37147 | GSM912291 | Duplicate |
| GSE37147 | GSM912296 | Duplicate |
| GSE37147 | GSM912305 | Duplicate |
| GSE37147 | GSM912342 | Duplicate |
| GSE37147 | GSM912273 | Duplicate |
| GSE37147 | GSM912305 | Duplicate |
| GSE37147 | GSM912342 | Duplicate |
| GSE37147 | GSM912342 | Duplicate |
| GSE37147 | GSM912348 | Duplicate |
| GSE37147 | GSM912376 | Duplicate |
| GSE37147 | GSM912376 | Duplicate |
| GSE37147 | GSM912376 | Duplicate |
| GSE37147 | GSM912463 | Poor Quality |
| GSE37147 | GSM912197 | Poor Quality |
| GSE37147 | GSM912300 | Poor Quality |
| GSE37199 | GSM913439 | Poor Quality |
| GSE37745 | GSM1019319 | Duplicate |
| GSE37745 | GSM1019246 | Duplicate |
| GSE37745 | GSM1019325 | Duplicate |
| GSE37745 | GSM1019247 | Duplicate |
| GSE37745 | GSM1019194 | Poor Quality |
| GSE37745 | GSM1019195 | Poor Quality |
| GSE37745 | GSM1019176 | Poor Quality |
| GSE37745 | GSM1019192 | Poor Quality |
| GSE37745 | GSM1019232 | Poor Quality |
| GSE37892 | GSM929512 | Poor Quality |
| GSE39491 | GSM970152 | Poor Quality |
| GSE39582 | GSM972249 | Duplicate |
| GSE39582 | GSM972472 | Duplicate |
| GSE39582 | GSM972243 | Duplicate |
| GSE39582 | GSM972044 | Duplicate |
| GSE39582 | GSM972091 | Duplicate |
| GSE39582 | GSM972090 | Duplicate |
| GSE39582 | GSM972245 | Duplicate |
| GSE39582 | GSM972473 | Duplicate |
| GSE39582 | GSM972515 | Duplicate |
| GSE39582 | GSM972248 | Duplicate |
| GSE43176 | GSM1057835 | Poor Quality |
| GSE46449 | GSM1130404 | Duplicate |
| GSE46449 | GSM1130406 | Duplicate |
| GSE46449 | GSM1130413 | Duplicate |
| GSE46449 | GSM1130417 | Duplicate |
| GSE46449 | GSM1130426 | Duplicate |
| GSE46449 | GSM1130428 | Duplicate |
| GSE46449 | GSM1130430 | Duplicate |
| GSE46449 | GSM1130434 | Duplicate |
| GSE46449 | GSM1130436 | Duplicate |
| GSE46449 | GSM1130468 | Duplicate |
| GSE46449 | GSM1130471 | Duplicate |
| GSE46449 | GSM1130483 | Duplicate |
| GSE48390 | GSM1176924 | Poor Quality |
| GSE48390 | GSM125120 | Poor Quality |
| GSE5462 | GSM125123 | Duplicate |
| GSE5462 | GSM125125 | Duplicate |
| GSE58697 | GSM1417097 | Poor Quality |
| GSE63885 | GSM1559328 | Duplicate |
| GSE63885 | GSM1559360 | Duplicate |
| GSE63885 | GSM1559385 | Duplicate |
| GSE63885 | GSM1559370 | Duplicate |
| GSE63885 | GSM1559375 | Duplicate |
| GSE63885 | GSM1559386 | Duplicate |
| GSE63885 | GSM1559361 | Poor Quality |
| GSE6532_U133PLUS2 | GSM151294 | Poor Quality |
| GSE6532_U133PLUS2 | GSM151280 | Poor Quality |

When transcriptomic data are processed in multiple batches, batch assignments can lead to confounding effects^23^. In the clinical annotations, we identified batch-processing information for datasets GSE25507, GSE37199, GSE39582, and GSE40292. We corrected for batch effects using the ComBat software^24^. The *Biomarker Benchmark* repository contains pre-and post-batch-corrected data. For dataset GSE37199, we identified two variables that could have been used for batch correction ("Centre" and "Plate"). Our repository contains batch-corrected data for both of these batch variables (the default is "Plate").

## Acknowledgements

SRP thanks Brigham Young University for research funds used in this study. AIB and PDH thank the BYU Office of Research and Creative Activities for research funds that supported this work. NPG thanks the Simmons Center for Cancer Research at Brigham Young University for a summer fellowship that supported this work. We thank researchers from many institutions who generated these data and released them to the public. We also thank the many research participants who participated in these studies.

## Author contributions

NPG: Collected data, wrote computer scripts, evaluated data quality, prepared figures and tables, wrote the manuscript.

AIB: Collected data, wrote computer scripts, evaluated data quality, edited the manuscript.

AB: Wrote computer scripts, wrote the manuscript.

PDH: Collected data, wrote computer scripts, edited the manuscript.

SRP: Collected data, wrote computer scripts, prepared figures and tables, wrote the manuscript.

